# Proteasome Caspase-Like Activity Regulates Stress Granules and Proteasome Condensates

**DOI:** 10.1101/2025.02.03.636236

**Authors:** Shirel Steinberger, Julia Adler, Nadav Myers, Yosef Shaul

## Abstract

The 20S proteasome maintains cellular protein homeostasis, particularly during stress responses. In a previous study, we identified numerous 20S proteasome substrates through mass spectrometry analysis of peptides generated from cellular extracts degraded by purified 20S proteasome. Many substrates were found to be components of liquid-phase separation, such as stress granules (SGs). Here, we demonstrate the degradation products predominantly arise from the caspase-like (CL) proteasomal activity. To investigate the functional implications of CL activity, we generated cell lines devoid of CL function by introducing the PSMB6 T35A mutation. These mutant cells exhibited slower growth rates, heightened sensitivity to stress, and activation of the unfolded protein response (UPR), as indicated by elevated levels of spliced XBP1 (sXBP1) and stress markers. Cells were subjected to arsenite and osmotic stress to assess their responses. Our findings reveal that CL activity is crucial for efficient SG assembly but does not significantly affect SG disassembly. Interestingly, in these mutant cells, proteasomes were more cytoplasmic under normal conditions but formed nuclear condensates/granules (PGs) upon NaCl osmotic stress. However, the PGs were unstable and rapidly disassembled. These findings underscore the important role of the proteasome’s CL activity in managing stress-induced dynamics of liquid-liquid phase, highlighting its importance in cellular adaptation to proteotoxic and genotoxic stress conditions.

## Introduction

The 26S proteasome is essential for cell survival and proliferation, playing a key role in regulating cellular protein quality control (Naujokat and Hoffmann 2002; Collins and Goldberg 2017; Coux, Tanaka, and Goldberg 1996). It removes damaged, misfolded, or unneeded proteins to maintain cellular homeostasis and prevent the accumulation of toxic aggregates (Pohl and Dikic 2019). This process is critical for various cellular functions, including cell cycle regulation, immune response, and stress adaptation. At the core of the 26S proteasome complex lies the 20S proteasome, which serves as its central component. The 20S proteasome is composed of four stacked rings, forming a cylindrical shape with a central channel where proteolysis occurs. These rings are made up of 28 protein subunits organized as follows: Two outer rings consist of seven α or PSMA subunits. Two inner rings consist of seven PSMB or β subunits (M Groll et al. 1997). The PSMB subunits form the proteolytic chamber, where the actual degradation of proteins takes place. Three of the PSMB subunits are catalytically active and possess protease activity, each with distinct specificities: PSMB6 (β1) exhibits CL (or peptidyl-glutamyl peptide hydrolyzing) activity, cleaving after acidic residues. PSMB7 (β2) exhibits trypsin-like activity, cleaving after basic residues. PSMB5 (β5) exhibits chymotrypsin-like activity, cleaving after hydrophobic residues (Kisselev, Akopian, Castillo, et al. 1999; Kim, Collins, and Goldberg 2018; M Groll et al. 1999). The chymotrypsin-like activity is generally considered the most crucial for its function. An important question is whether each of the three types of proteolytic activities of the 20S catalytic particle has a unique role in these processes. Here, we addressed this question by generating cell lines devoid of CL activity.

We investigated the role of CL activity in the formation of stress granules (SGs) and proteasome granules. SGs are cytoplasmic, membraneless organelles formed via liquid–liquid phase separation (LLPS) in response to stressors like heat shock, oxidative stress, or nutrient deprivation, driven by interactions involving RNA-binding proteins like G3BP (Protter and Parker 2016) (Guillén–Boixet et al. 2020; Yang et al. 2020; Gwon et al. 2021). Containing untranslated mRNAs, RNA-binding proteins, translation factors, and signaling components, SGs regulate RNA metabolism and stress signaling (Hofmann et al. 2021; Ries et al. 2019). Their formation is primarily controlled by eIF2α phosphorylation and RNA-binding proteins, which mediate nucleation and stabilization (N. L. Kedersha et al. 1999; Millar et al. 2023).

SG dynamics depend on the balance between aggregation and disaggregation, influenced by chaperones (e.g., Hsp70, Hsp90) (Mediani et al. 2021), post-translational modifications (e.g., phosphorylation, ubiquitination, methylation) (Z. Wang et al. 2023), and remodeling enzymes (e.g., USP10, DDX3) (Buchan 2024). Disassembly, driven by eIF2α dephosphorylation, translation restoration, and disaggregase activity, occurs when stress resolves, restoring normal cellular functions. Dysregulated SG dynamics are linked to diseases like cancer, viral infections, and neurodegeneration (Li et al. 2013; Markmiller et al. 2018).

The role of proteasomes in SG assembly and dissolution is not well understood. Inhibition of the Ubiquitin-Proteasome System (UPS) Induces SGs Formation (Mazroui et al. 2007). ZFAND1 is a regulator of SG clearance, that interacts with the 26S proteasome and p97 (valosin containing protein (VCP), a chaperone like protein), which are recruited to arsenite-induced SGs for their dissolution (Turakhiya et al. 2018). Proteasome inhibitors also indirectly might regulate SG formation through certain cellular pathways, mRNA metabolism, cellular stress responses, and protein quality control systems. However, whether and which of the proteasomal different proteolytic activities are directly involved in SG formation and dissolution awaits further investigation.

Under certain conditions, proteasome condensates, referred to as proteasome granules (PGs), exhibiting properties of liquid-liquid phase separation (LLPS), are formed preferentially in the nuclei of the animal cells (Yasuda et al. 2020; Steinberger, Adler, and Shaul 2023). Some of the essential components of PGs are p97 (VCP), RAD23B, a substrate-shuttling factor for the proteasome, and UBE3A, ubiquitin-protein ligase E3A also known as E6AP (Yasuda et al. 2020). Mechanistically, LLPS assembly is mediated via multivalent interactions of two ubiquitin-associated domains of RAD23B and ubiquitin chains consisting of four or more ubiquitin molecules (Yasuda et al. 2020).

In yeast, proteasome condensate, also known as proteasome storage granules (PSGs), are formed under certain stress conditions when reaching stationary phase growth or nutrient deprivation (Laporte et al. 2008; Gu et al. 2017; Waite et al. 2024). Unlike the nuclear PG in animal cells, the PSGs are cytosolic (Laporte et al. 2008). Mechanistically, PSGs contain ubiquitin chains together with Rad23 and Dsk2, the proteasome shuttle factors, which are critical component for PSG formation (Waite et al. 2024), highlighting the mechanistic similarity between PGs and PSGs.

When the cells were treated with the proteasome inhibitor MG-132, an inhibitor of chymotrypsin-like activity, or b-AP15, a specific inhibitor of the deubiquitinating enzymes, the number and size of the PGs increased, and the rate of clearance was significantly delayed (Yasuda et al. 2020). However, the importance of proteasome CL activity in PG formation remained an open question. Here, we report the generation of cell lines devoid of CL activity and demonstrate that they are inferior in the level of SGs and PGs formation under stress.

## Materials and Methods

### Materials

Sodium (meta)arsenite, doxorubicin, antimycin A, rotenone, poly-D-lysine hydrobromide were purchased from Sigma-Aldrich (Burlington, MA, USA); Hoechst33342 from Molecular Probes (Eugene, OE, USA); bortezomib from APExBIO (Houston, TX, USA). Cisplatin was from ABIC (Israel).

### Cells and Cell Culture

PSMB6 T35A HEK293 mutant cell generation with reduced or abolished proteasome CL activity is described (Reuven et al. 2021). To introduce the YFP tag to PSMB6 and PSMB6 T35A genes, we used CRISPR technology, as previously described (Reuven et al. 2019, 2021). Cell lines were also labeled with the stress granules marker, G3BP1-mCherry, as described elsewhere (Steinberger, Adler, and Shaul 2023). Cells were grown at 37 °C in a humidified incubator with 5.6% CO2 in Dulbecco’s modified Eagle’s medium (DMEM; GIBCO, Life Technologies, Thermo Scientific, Waltham, MA, USA) supplemented with 8% fetal bovine serum (GIBCO), 100 units/mL penicillin, and 100 μg/mL streptomycin (Biological Industries, Beit Hemek, Israel). The Incucyte^®^ SX1 live-cell analysis system (Sartorius) was used to photograph cells and quantify cell proliferation and viability.

### Live Cell Imaging

Cells were seeded on a 96-well glass bottom Microwell plate 630 µL black 17 mm low glass from Matrical Bioscience (MGB096-1-2-LG), coated with poly-D-lysine hydrobromide, and were allowed to adhere overnight. Before the imaging, cell media were supplemented with five µM Hoechst 33342. The plate was placed in the microscope chamber, and cells were maintained at 37 °C and 5% CO_2_ for the duration of the experiment. Osmotic stress was induced by replacing the media at time zero with media supplemented with 150 mM NaCl or 200 mM sucrose. For the induction of proteotoxic stress, the medium was replaced with one mM of sodium (meta)arsenite. Images of four different sites in a well were taken every 4 min for three hours using a VisiScope Confocal Cell Explorer live cell imaging system with a 60x-oil objective. Data was processed and analyzed as described in (Steinberger, Adler, and Shaul 2023). Statistical tests of the two-tailed t-test were performed to assess significance using Excel.

### Immunoblot Analysis

SDS-PAGE and immunoblotting were performed as previously described (Levy et al. 2007) using RIPA buffer (50 mM Tris-HCl pH 7.5, 150 mM NaCl, 1% Nonidet P-40 (v/v), 0.5% deoxycholate (v/v), 0.1% SDS (w/v)) supplemented with a cocktail of protease inhibitors (APExBIO, Houston, TX) for cell extract preparation. Antibodies used in this study: pS52 eIF2α (Thermo Scientific, Rockford, Il, USA), ATF4 (Cell Signaling, Beverly, MA, USA), eIF2α, and hsp90 (Santa Cruz Biotechnology, Santa Cruz, CA, USA). Horseradish peroxidase-conjugated secondary antibodies were from Jackson ImmunoResearch Laboratories, West Grove, PA. Enhanced chemiluminescence was performed with the EZ-ECL kit (Biological Industries, Beit Hemek, Israel), and signals were detected by the ImageQuant LAS 4000 (GE Healthcare, Piscataway, NJ, USA).

#### 20S Proteasome degradation

Dataset used for analysis - PXD010132 from the PRIDE (*), https://www.ebi.ac.uk/pride/, MS proteomics repository described in (Myers et al. 2018).

### Protein Analysis

20S Proteasome degradation: Dataset used for analysis - PXD010132 from the PRIDE (Vizcaíno et al. 2016), https://www.ebi.ac.uk/pride/, MS proteomics repository described in (Myers et al. 2018).

All protein sequences were retrieved from the UniProt database (http://www.uniprot.org/) using the curated Swiss-Prot knowledgebase (Consortium, 2012). Statistical analyses and data visualization were conducted using MATLAB 2016b (The MathWorks, Natick, 2014). For hypothesis testing, p-values were calculated using a two-sided non-parametric Wilcoxon rank-sum test applied to continuous data. While all data points were included in the statistical analyses, outliers were excluded from box plots to ensure clarity and improve visualization.

To identify amino acid preferences around the cleavage site (P5–P5’) in 20S core particle (CP) substrates, frequency profiles were generated using IceLogo (Colaert et al. 2009). This tool highlights statistically over- or under-represented amino acids, providing insights into sequence motifs associated with proteolytic cleavage.

### RNA extraction, cDNA preparation and analysis

Total RNA was extracted using TRI Reagent (MRC, Beverly Hills, CA, USA). First-strand cDNA synthesis was performed using the iScript cDNA Synthesis Kit (Quanta, Houston, TX, USA). Quantitative real-time PCR (qRT-PCR) was conducted using the LightCycler 480 (Roche, Basel, Switzerland) with PerfeCta^®^ SYBR Green FastMix (Quanta). All qPCR results were normalized to TBP1 mRNA levels.

### 2.4. Statistical Analysis

The statistical significance (*p*-value) between means was assessed by two-tailed Student’s *t*-tests.

## Results

### Caspase-like activity dominates substrate degradation by 20S proteasome in vitro

In previous experiments, we incubated heat-stable proteins derived from cell extracts enriched for intrinsically disordered proteins and protein regions (IDPs/IDRs) with purified 20S proteasomes to identify potential 20S substrates (Myers et al. 2018). Proteasomes are known for their length-specific degradation, predominantly generating oligopeptides ranging from 3 to 25 amino acid residues (Kisselev, Akopian, Woo, et al. 1999; Voges, Zwickl, and Baumeister 1999; Michael Groll and Clausen 2003). We compared the sizes of the oligopeptides produced by the 20S proteasome with those generated by trypsin digestion of the heat-stable proteins. Notably, the 20S proteasome produced shorter peptides compared to trypsin (Fig. 1A). Further analysis of the mass spectrometry (MS) data revealed that the peptides generated by the 20S proteasome spanned the entire length of the substrate proteins. This suggests that the 20S proteasome does not exhibit a specific preference for cleavage sites, as the degradation occurs uniformly across the substrate (Fig. 1B).

**Figure 1.**
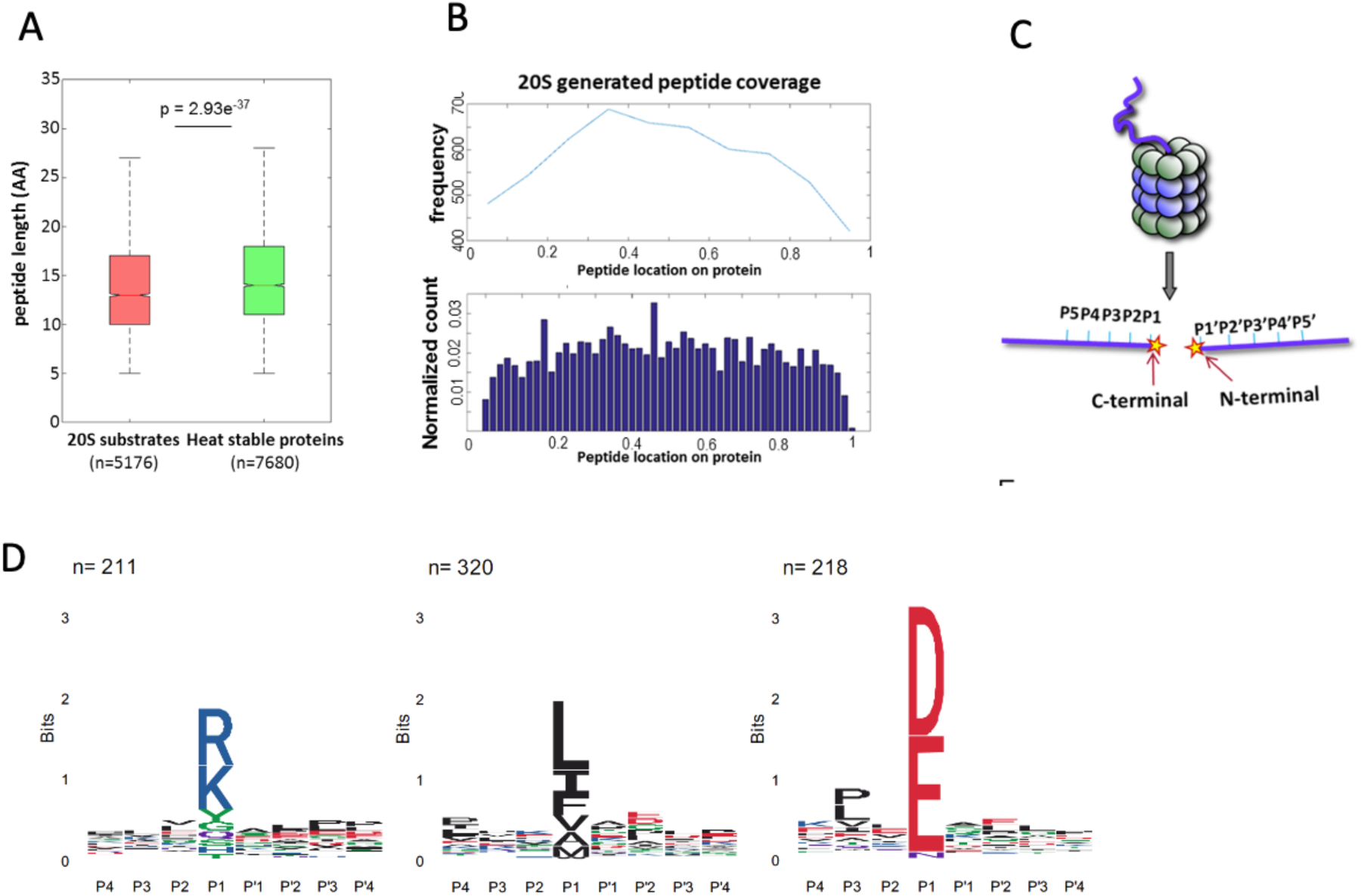
Characterization of 20S proteasome substrates and cleavage specificity. (A) Box plot comparing the peptide lengths of 20S proteasome-generated products (red, n=5176) with those derived from trypsin digestion of heat-stable proteins (green, n=7680). The 20S proteasome generates shorter peptides on average (p = 2.93e−37). (B) Distribution of peptide locations across substrate proteins as determined by mass spectrometry. The upper panel shows the frequency of peptides spanning the entire substrate, while the lower panel displays the normalized count of peptide locations, indicating uniform degradation without specific cleavage preferences. (C) Schematic representation of substrate positioning within the proteolytic chamber of the 20S proteasome. The regions adjacent to the cleavage site, P5–P1 on the C-terminal side and P1’–P5’ on the N-terminal side, influence cleavage efficiency and specificity. (D) Sequence preferences of the 20S proteasome cleavage sites, illustrated by IceLogo analysis. Peptides digested by proteasomes in vitro clustered to acidic, basic, and hydrophobic residues in the cutting site. Notably, acidic residues (D, E) are enriched at the P1 position, consistent with CL activity under experimental conditions.

The structural architecture of the 20S proteasome proteolytic chamber plays a crucial role in substrate specificity. Binding pockets surrounding the active sites determine substrate affinity, guiding the interaction between substrates and the proteasome active sites. Consequently, the amino acid sequence around the scissile bond significantly influences degradation efficiency and specificity (Michael Groll and Clausen 2003; Huber et al. 2012). The amino acid positions immediately preceding and following the cleavage site are illustrated (Fig. 1C). To gain a deeper understanding of the sequence preferences of 20S proteasome substrates, we analyzed the amino acid composition at the N-terminal (P1’–P5’) and C-terminal (P5–P1) regions of the peptides generated by the 20S proteasome. This analysis was performed using our peptide database derived from 20S proteasome-generated products (Myers et al. 2018). The IceLogo software (Colaert et al. 2009) was used to generate a frequency profile that highlights amino acid preferences or disfavor surrounding the cleavage site (P5-P5’) in 20S proteasome substrates. This analysis reveals a distinct degradation sequence characteristic of 20S proteasome activity (Fig. 1D). Notably, the enrichment of acidic residues at the P1 position indicates a strong preference for CL activity within the proteasome under the tested conditions. These findings suggest that the 20S proteasome CL activity plays a dominant role in the degradation of the IDPs/IDRs enriched substrates.

### Proteasome CL activity deficiency impairs cell growth, disrupts stress responses, and activates the unfolded protein response (UPR)

To evaluate the role of CL activity in the cells, we generated both heterozygote and homozygote PSMB6 T35A edited cell lines, demonstrating partial or lack of CL activity, respectively, as previously described (Reuven et al. 2021). The growth rate of the mutants was much slower, especially in the PSMB6 T35A homozygote cells (Fig. 2 A and B). Next, we treated the cells with bortezomib, an inhibitor of proteasome chymotrypsin-like activity, and the growth of homozygote cells was severely compromised (fig 2C). The edited cells show a much lower growth rate when treated with cisplatin, a DNA damage agent (fig 2D), and doxorubicin, a DNA intercalating agent (fig 2E). Inhibition of cellular respiratory by each Antimycin A and rotenone reduced the growth of the PSMB6 T35A edited cells (fig 2F and G). Given the important role of proteasome in reducing ER stress and unfolded protein response (UPR), we asked whether cells deficient in CL activity induce the UPR. To this end, the levels of some of the UPR markers were measured (Read and Schröder 2021). The level of spliced XBP-1 (sXBP-1) mRNA was increased in the homozygote PSMB6 T35A edited cells (fig 2H). ATF4 and p-eIF2*α* levels were analyzed by immunoblot and showed significant induction in the PSMB6 T35A edited cells (fig 2I). The T35A mutation in PSMB6, generating proteasomes deficient in CL activity, appears to enhance the activation of the UPR, as evidenced by increased spliced XBP1 mRNA and elevated stress markers. These data demonstrate that PSMB6 T35A mutation, impairing CL activity, slows cell growth and sensitizes cells to proteotoxic and genotoxic stress. The mutation also induces UPR activation.

**Figure 2.**
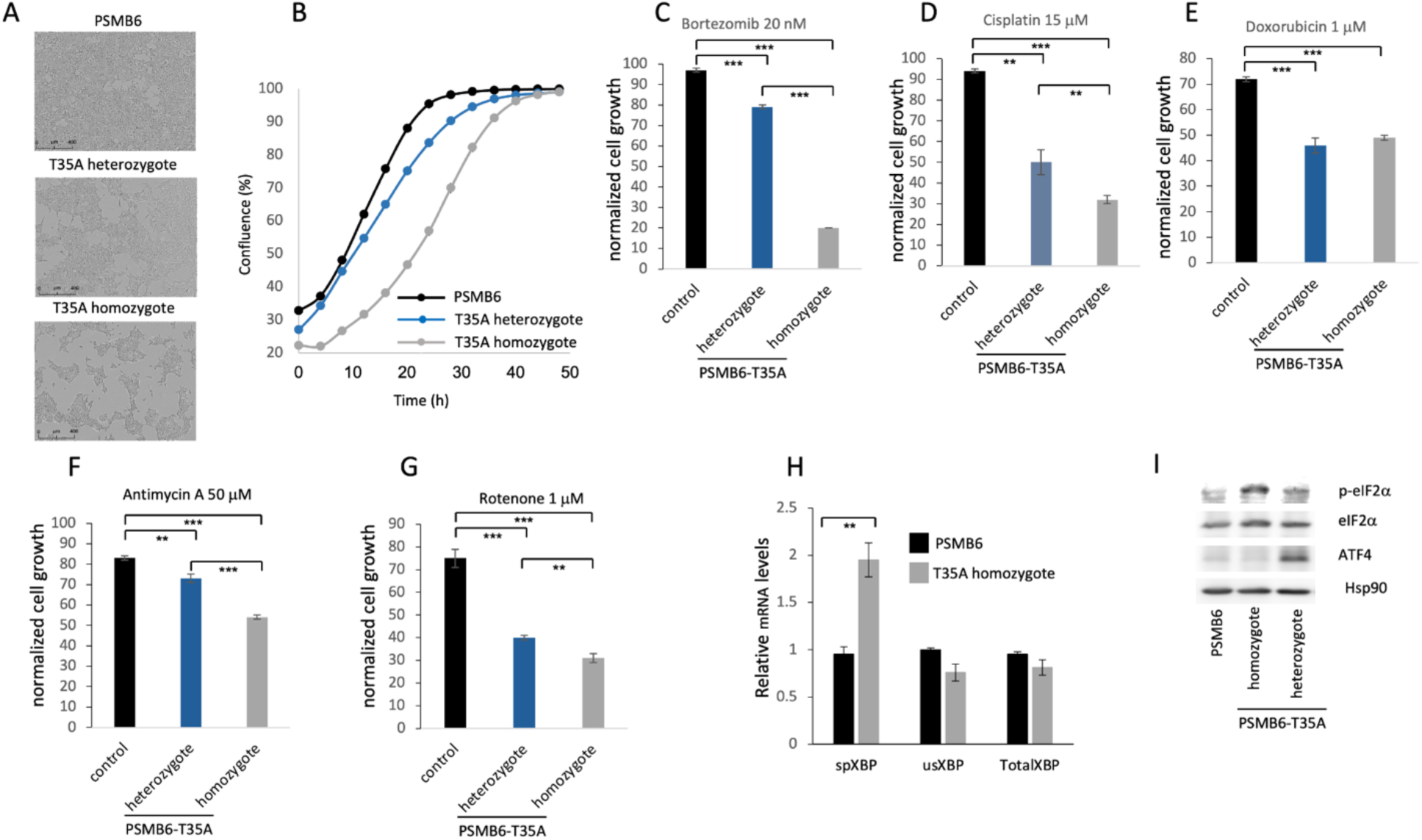
Effects of PSMB6 T35A mutation on cell growth, stress response, and unfolded protein response (UPR). (A) Representative phase-contrast images of wild-type PSMB6, heterozygous, and homozygous PSMB6 T35A mutant cells, showing slower growth in the mutants. (B) Growth curves of PSMB6 wild-type, heterozygous, and homozygous T35A mutant cells over 50 hours, using Incucyte® SX1 live-cell analysis system. A significantly reduced rate of confluence achievement in mutant cells, particularly in homozygotes, is shown. (C-E) Growth rates of PSMB6 T35A mutant and wild-type cells treated with bortezomib, an inhibitor of proteasome chymotrypsin-like activity (C), cisplatin, a DNA-damaging agent (D), or doxorubicin, a DNA-intercalating agent (E). Homozygous mutants show severe growth inhibition upon treatment. (F-G) Effect of mitochondrial inhibitors, antimycin A (F) and rotenone (G), on cell growth. (H) Relative levels of spliced XBP-1 (sXBP-1), unspliced XBP-1 (usXBP-1), and total XBP-1 mRNA were measured by qRT-PCR. Spliced XBP-1 levels are elevated in homozygous PSMB6 T35A cells, indicating UPR activation. (I) Immunoblot analysis of UPR markers (p-eIF2α and ATF4) in PSMB6 wild-type and T35A mutants. Homozygous PSMB6 T35A cells exhibit elevated levels of p-eIF2α and ATF4. Hsp90 is used as a loading control to normalize protein expression. ** *p* < 0.01, *** *p* < 0.001.

### Proteasome caspase-like activity regulates stress granules

Previously, we reported that the purified 20S proteasome degrades proteins associated with stress granules (Myers et al. 2018). As shown above (figure 1D), CL activity was the primary factor in degrading these proteins. This prompted us to investigate whether proteasome CL activity regulates stress granule (SG) assembly. To this end, cells expressing G3BP-mCherry, a marker of SGs, were treated with arsenite, a well-known inducer of SG formation. Fluorescent imaging over time (0–176 minutes) showed robust SG assembly in PSMB6 wild-type cells, with the number of SGs gradually increasing up to 88-120 minutes before slowly disassembling (Fig. 3A). By contrast, PSMB6 T35A mutant cells, devoid of CL activity, exhibited significantly fewer SGs per cell (Fig. 3A, B). Interestingly, despite the reduced number of SGs in mutant cells, their size was comparable to those in wild-type cells (Fig. 3C).

**Fig 3:**
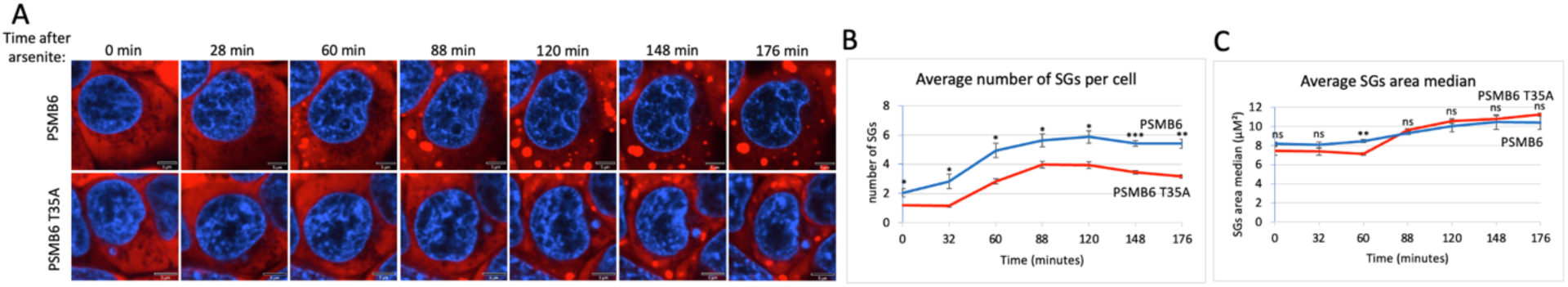
SGs dynamic over time after arsenite treatment in wild-type and PSMB6 T35A mutant cells. (A) Shows fluorescent images of cells at various time points (0 to 176 minutes) after arsenite treatment. Blue represents nuclear staining (DAPI). Red represents G3BP-mCherry expression. The scale bar is 5µm. In PSMB6 wild-type cells, SG formation appears robust and increases over time after arsenite treatment. In PSMB6 T35A mutant cells, SG formation is reduced or delayed, with fewer SGs evident compared to the wild-type cells. (B) The average number of stress granules per cell over time. Note that SGs form rapidly in PSMB6 wild-type cells, reaching a peak between 88 and 148 minutes. Meanwhile, in PSMB6 T35A mutant cells, SG formation is significantly reduced at most time points, as indicated by statistical markers. (C) SG area median (µm²) per cell over time. The data were analyzed by 2-tailed unpaired t-test * *p* < 0.05, ** *p* < 0.01, *** *p* < 0.001.

Quantification revealed that wild-type cells rapidly formed SGs, peaking between 88 and 148 minutes post-treatment, while T35A mutant cells showed consistently reduced SG formation. The median SG area (µm²) per cell was similar between the two cell types, suggesting that CL activity primarily impacts SG number rather than size. The relatively parallel curves for SG disassembly in both cell types indicate that the disassembly rate is similar, regardless of CL activity. These findings suggest that proteasome CL activity is crucial for efficient SG assembly but does not significantly affect SG disassembly. Together, these results highlight the proteasome’s critical role in regulating SG dynamics during arsenite-induced proteotoxic and oxidative stress responses.

### Proteasome caspase-like activity regulates SG dynamics under osmotic stress

Osmotic stress, typically induced by high concentrations of NaCl or other osmolytes, effectively triggers the formation of SGs (Protter and Parker 2016; Hu et al. 2023). To explore the role of CL activity in SG assembly during osmotic stress, we treated cells with NaCl. Unlike arsenite treatment, NaCl-induced SG assembly occurred much faster, reaching its maximal level within 33 minutes. However, the number of SGs per cell was significantly lower in PSMB6 T35A mutant cells compared to wild-type cells (Fig. 4A and B). Interestingly, the size of SGs in the mutant cells increased over time and became significantly larger than those in wild-type cells (Fig. 4C). Despite these differences, the disassembly rate was similar between wild-type and mutant cells, as indicated by the parallel behavior of the disassembly curves. Next, we induced osmotic stress by treating cells with sucrose. Under this condition, SG formation was slower in PSMB6 T35A mutant cells, although the disassembly rate remained comparable to wild-type cells (Fig. 4D and E). The total number of SGs formed was significantly lower in PSMB6 T35A cells (Fig. 4E), and the size of SGs was smaller at the early time but increased later on (Fig. 4F). These findings suggest that proteasome CL activity is critical for the efficient assembly of SGs under both proteotoxic arsenite stress and osmotic stress conditions. The data highlight the role of CL activity in regulating SG formation, particularly under NaCl-induced stress.

**Fig 4:**
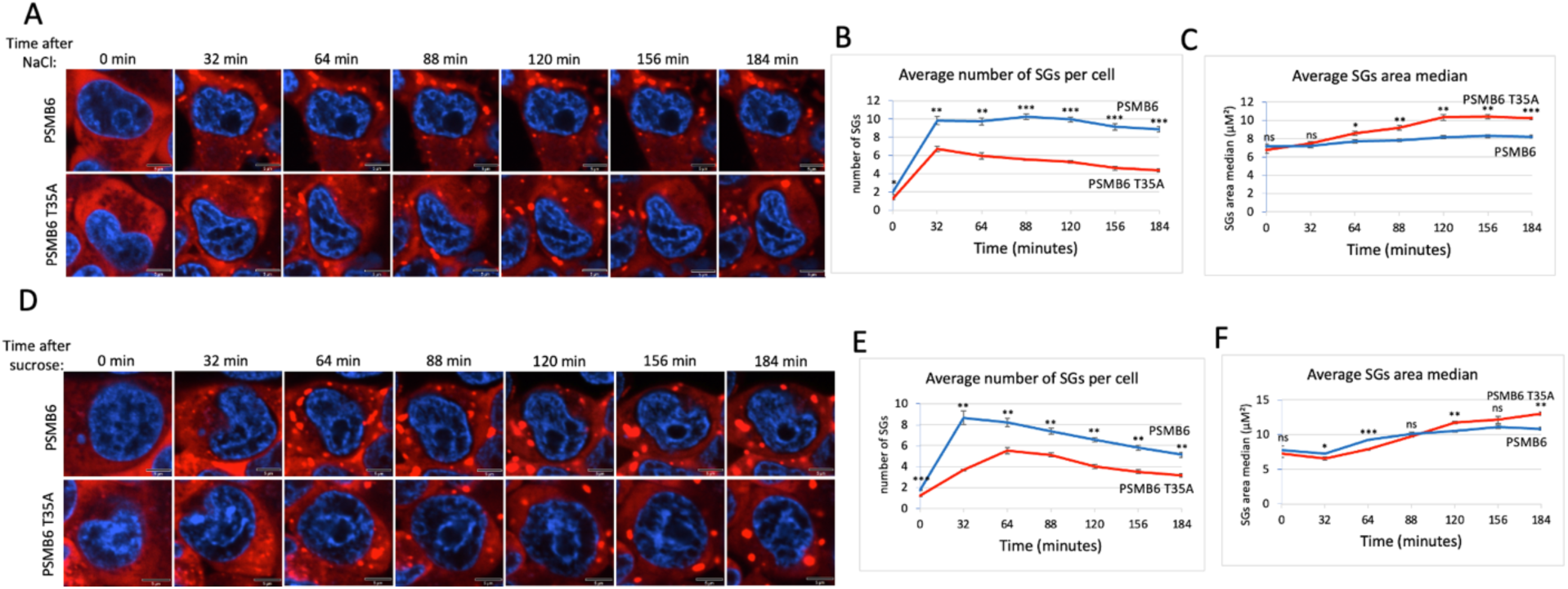
SG dynamic over time in wild-type and PSMB6 T35A mutant cells after osmotic stress. (A) Microscopic images of wild-type G3BP-mCherry HEK293 (PSMB6) and PSMB6 T35A mutant cells at the indicated time points. Blue represents nuclear staining (DAPI). Red represents G3BP-mCherry expression. Stained for stress granules (red) and nuclei (blue) at different time points. SGs are increased after NaCl treatment in both cell types, but their formation appears more pronounced or persists longer in the wild-type condition compared to the mutant. (B) The average number of stress granules per cell at different time points. The blue line (wild-type) shows a higher and sustained number of SGs over time. The red line (mutant) shows a lower peak and faster reduction in SG numbers. (C) As in panel B but the average SGs area median over time for both cell types shown. (D) Similar to panel A but cells were treated with sucrose. (E) Similar to panel B but cells were treated with sucrose. (F) Similar to panel C but cells were treated with sucrose. The data were analyzed by 2-tailed unpaired t-test * *p* < 0.05, ** *p* < 0.01, *** *p* < 0.001.

### Proteasome caspase-like activity regulates proteasome granule dynamic

It has been reported that under osmotic stress, proteasomes rapidly form nuclear granules (condensates) in cells (Yasuda et al. 2020; Steinberger, Adler, and Shaul 2023). Inhibition of the proteasome chemotrypsin-like activity did not affect the process (Yasuda et al. 2020). Therefore, we asked whether CL activity is required to assemble proteasome granules. To address this, we used CRISPR technology to YFP tag the PSMB6 and PSMB6 T35A genes, as previously described (Reuven et al. 2019; Steinberger, Adler, and Shaul 2023). Interestingly, we observed that proteasomes in PSMB6 T35A cells were significantly less localized in the nucleus and showed higher levels in the cytoplasm (Fig. 5A). Quantification confirmed a significant reduction in the nuclear-to-cytoplasmic intensity ratio in PSMB6 T35A cells compared to wild-type PSMB6 cells (Fig. 5B).

**Figure 5.**
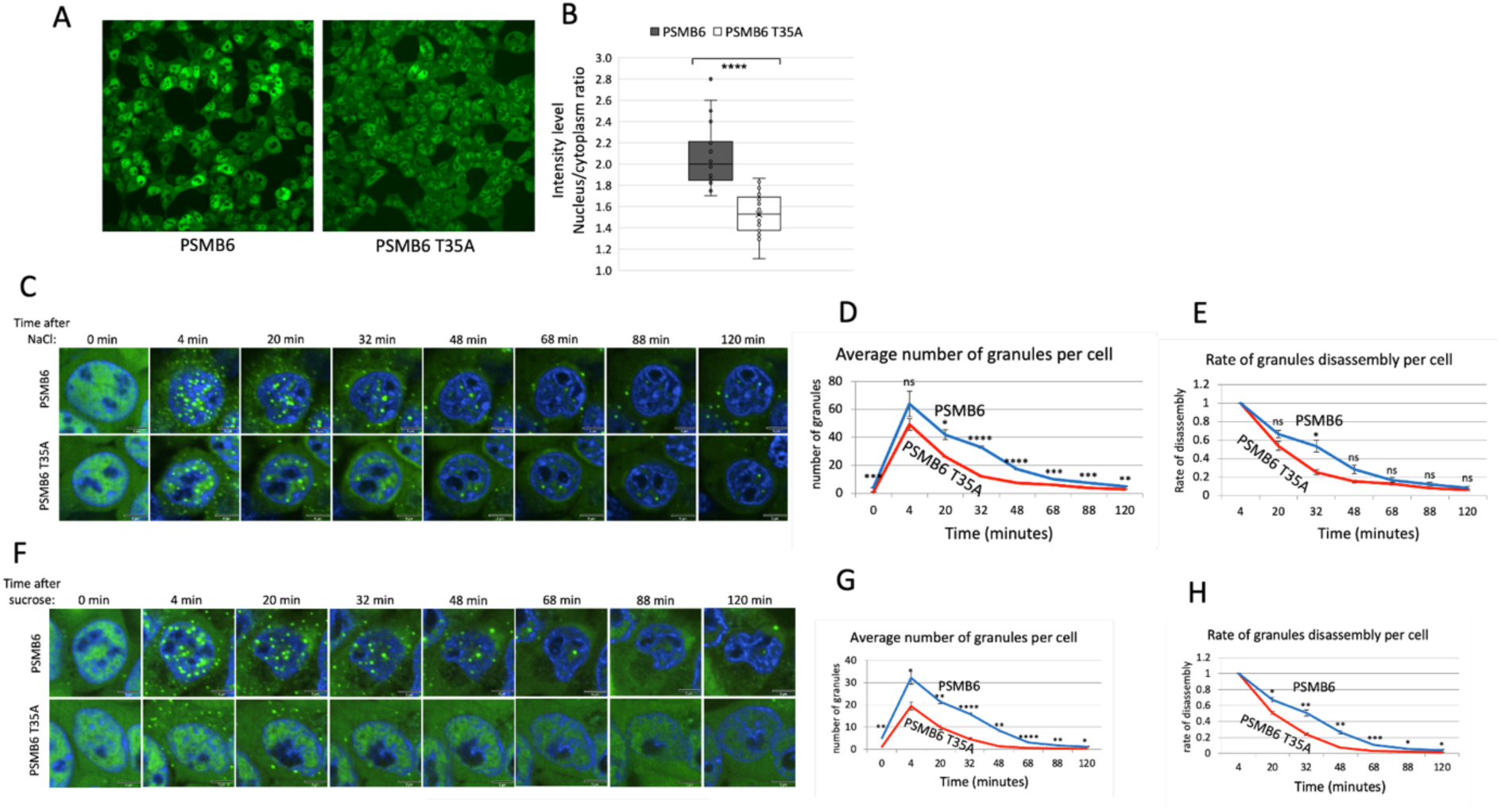
PSMB6 T35A cells exhibit altered nuclear localization and rapid disassembly of PGs under osmotic stress. (A) Representative images showing subcellular localization of proteasomes in PSMB6 and PSMB6 T35A cells under normal conditions. YFP-tagged PSMB6 displays strong nuclear localization, while PSMB6 T35A exhibits reduced nuclear localization and increased cytoplasmic distribution. (B) Quantification of nuclear-to-cytoplasmic intensity ratios of YFP-tagged proteasomes in PSMB6 and PSMB6 T35A cells. Data indicate a significant reduction in nuclear localization in PSMB6 T35A cells (****p < 0.0001). (C, F) Time-lapse images of PG formation and disassembly following osmotic stress induced by NaCl (C) or sucrose (F). Both PSMB6 and PSMB6 T35A cells form PGs within four minutes of treatment. (D, G) Quantification of the average number of PGs per cell over time after NaCl (D) or sucrose (G) treatment. PSMB6 T35A cells exhibit significantly fewer PGs at their peak and a faster rate of disassembly compared to PSMB6 cells. (E, H) Rate of PG disassembly per cell following NaCl (E) or sucrose (H) treatment. PSMB6 T35A cells show a significantly faster disassembly rate (**p < 0.01, ***p < 0.001).

Next, we exposed the cells to osmotic stress using NaCl. As expected, we observed the formation of proteasome granules (PGs) within four minutes in both PSMB6 and PSMB6 T35A cells (Fig. 5C). However, the number of PGs per cell reached a lower peak in PSMB6 T35A cells. It disassembled more rapidly than PSMB6 cells (Fig. 5D and 5E), suggesting that the mutation impacts the stability of PGs under stress.

Similar results were observed when the cells were subjected to osmotic stress induced by sucrose (figs. 5F–5H). In this case, the maximum number of PGs per cell in PSMB6 T35A cells was significantly lower than in wild-type cells. Additionally, the PGs in PSMB6 T35A cells were predominantly localized in the cytoplasm. This cytoplasmic localization and the rapid disassembly of PGs further support the notion that the CL activity is not essential for the initial formation of the PGs but is crucial for maintaining their stability. This highlights a potential role for CL activity in regulating proteasome condensate dynamics under stress conditions.

### PSMB6 T35A mutant cells enhance arsenite-induced proteasome granules

The formation of PGs in animal cells in response to osmotic stress has been previously documented (Yasuda et al. 2020; Steinberger, Adler, and Shaul 2023), however, whether proteotoxic stress induces the formation of PGs has not been examined in depth. Earlier studies reported that arsenite does not induce the PGs up to 60 minutes of treatment, whereas osmotic stresses initiate their formation within a few minutes (Yasuda et al. 2020; Steinberger, Adler, and Shaul 2023). In this study, we treated cells with arsenite for a longer duration and revealed that PGs formed after 80 minutes of treatment. Surprisingly, the PGs are not nuclear but exclusively localized in the cytoplasm (Fig. 6A). Interestingly, in arsenite-treated PSMB6 T35A cells, discrete punctate structures, likely PGs, form much earlier-60 minutes after treatment. The behavior of the proteasome under arsenite treatment differs from that observed with osmotic stress in terms of cytosolic localization and facilitated assembly.

**Figure 6.**
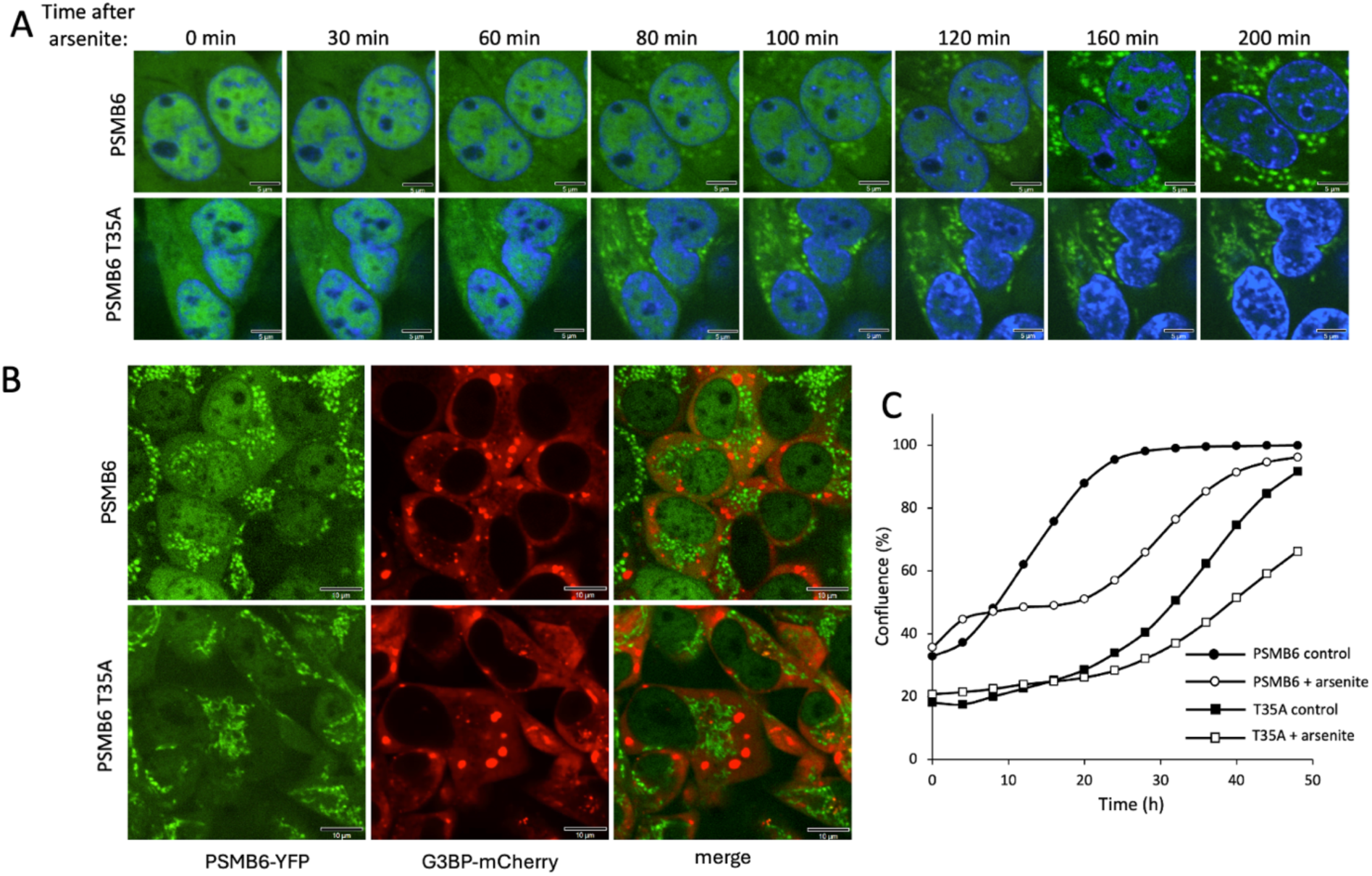
Proteasome granule formation and cell growth dynamics under arsenite-induced stress. (A) Time-course analysis of proteasome granule formation in PSMB6 and PSMB6 T35A cells following 3hrs treatment of 1 mM arsenite. PGs (green) appear in the cytoplasm after 80 minutes in PSMB6 cells and after 60 minutes in PSMB6 T35A cells, as observed by fluorescence microscopy. Nuclei are stained with DAPI (blue). Scale bars, 5 µm. (B) Simultaneous analysis of PGs (PSMB6-YFP, green) and SGs (G3BP-mCherry, red) in PSMB6 and PSMB6 T35A cells under arsenite treatment. No colocalization is observed, indicating that proteasome and stress granules are distinct entities. Scale bars, 10 µm. (C) Growth kinetics of PSMB6 and PSMB6 T35A cells under control and arsenite-treated conditions. Arsenite-treated PSMB6 cells show biphasic growth with recovery after 20 hours, whereas arsenite-treated PSMB6 T35A cells exhibit slower growth and fail to recover by 50 hours. The growth curve was analyzed *in situ* using the Incucyte® SX1 live-cell analysis system.

The kinetics of PGs formation under arsenite treatment resembles those of SGs based on their time of appearance and cytoplasmic localization. To investigate whether PGs are components of stress granules, we simultaneously analyzed cells for the two types of granules. No colocalization was observed, suggesting that proteasome and stress granules are distinct entities (Fig. 6B). This distinction was consistent in both PSMB6 and PSMB6 T35A cells.

Next, we examined the growth dynamics of edited and arsenite-treated cells. Arsenite-treated PSMB6 cells exhibited biphasic growth: their growth rate was slower than untreated cells initially but recovered after 20 hours to match the growth rate of untreated cells (Fig. 6C). In contrast, untreated PSMB6 T35A cells displayed a slower growth rate than unedited cells but eventually reached confluence after 50 hours. However, arsenite-treated PSMB6 T35A cells grew significantly slower and failed to recover after 50 hours. These data suggest that arsenite induces cytosolic PGs and that PSMB6 T35A cells are more susceptible to arsenite treatment, exhibiting earlier granule formation and slower cell growth.

## Discussion

This study highlights the role of the 20S proteasome’s caspase-like (CL) activity in cellular stress responses. Our findings show that cells devoid of CL activity increase cellular sensitivity to stress, resulting in slower proliferation and heightened vulnerability to arsenite and NaCl-induced stress. The unfolded protein response (UPR) activation in CL-deficient cells, marked by elevated sXBP-1 and stress markers, suggests that CL activity mitigates proteotoxic and genotoxic stress by preventing the accumulation of misfolded or aggregated proteins. Using in vitro 20S proteasome reactions and proteomic approaches, we show that CL activity selectively degrades intrinsically disordered proteins (IDPs) and regions (IDRs) (Myers et al. 2018), which are key components of liquid-liquid phase separation (J. Wang et al. 2018), such as SGs and proteasome condensates. Cells harboring this mutation exhibited slower SG assembly during arsenite-induced proteotoxic stress and reduced SG stability under osmotic stress. These observations highlight the dual role of CL activity in facilitating both the efficient assembly and the structural integrity of SGs. Furthermore, the rapid disassembly of proteasome condensates in CL-deficient cells underscores the importance of CL activity in sustaining the dynamics of these assemblies under stress conditions.

The delay in SG assembly observed in cells lacking CL activity raises intriguing questions about the molecular mechanisms involved. Previous studies proposed that SG assembly is driven by interactions among RNA molecules, SG proteins like G3BP1, and homotypic/heterotypic interactions involving IDRs (N. Kedersha et al. 2016; Panas, Ivanov, and Anderson 2016; Jain and Vale 2017; Van Treeck and Parker 2018; Protter et al. 2018). More recently, it has been reported that RNA binding disrupts an autoinhibitory interaction within G3BP1, facilitating SG assembly (Guillén–Boixet et al. 2020). Our findings suggest a potential regulatory role for CL activity in SG dynamics, either through direct processing of G3BP1 to relieve autoinhibition or indirectly via peptides generated during proteasomal degradation.

Interestingly, while CL activity was not strictly required for the initial formation of proteasome granules under osmotic stress, mutant cells displayed fewer and less stable granules. This unexpected result suggests a potential indirect role for CL activity, linked to stress-induced changes in CL-deficient cells. Alternatively, it raises the intriguing possibility of a direct role for CL activity in granule formation, perhaps by processing essential components or regulators. Additionally, the acidic peptides generated by CL activity may directly contribute to proteasome granule formation, warranting further investigation.

Under inflammation, animal cells express three proteasomal proteolytically active subunits: β1i, β2i, and β5i, collectively termed immunosubunits due to their cytokine-inducible expression and role in antigenic peptide generation (Schmidt and Finley 2014; Nathan et al. 2013; Seifert et al. 2010). These subunits replace the constitutive proteasome subunits β1, β2, and β5. β1i provides chymotrypsin-like activity, replacing the caspase-like activity of β1, rendering the immunoproteasome’s caspase-like activity nearly nonfunctional. Whether the unique behavior of cells lacking caspase-like activity is relevant under inflammation signaling remains an open question.

Our findings highlight the multifaceted role of the 20S proteasome’s CL activity in maintaining proteostasis under stress conditions. Possibly by targeting IDP/IDR-enriched substrates, this activity regulates SG dynamics and proteasome granules stability, processes essential for cellular adaptation to stress (Li et al. 2013; Markmiller et al. 2018; N. Kedersha and Anderson 2009). Future studies should focus on elucidating the molecular mechanisms underlying the selective targeting by CL activity and its broader implications in diseases associated with proteostasis imbalances, such as neurodegenerative disorders and cancer.

